# Lysosomal Ion Homeostasis Drives Delayed Hair Cell Death After Aminoglycoside Uptake

**DOI:** 10.64898/2026.06.10.731244

**Authors:** Francisco Barros-Becker, Ananya Cholkar, Tor H. Linbo, David W. Raible

## Abstract

Aminoglycoside ototoxicity has been widely reported and remains an important public health issue. Unfortunately, the molecular mechanisms of ototoxicity are not well understood. Here, we report the lysosome compartment as the main driver of delayed cell death triggered by aminoglycosides. By labeling early and late endosomes we show that endocytosis is not an significant path of aminoglycoside uptake. Instead, we show that aminoglycosides are delivered to lysosomes primarily through autophagy. Hair cells can be protected from damage by activation of the dual function lysosomal Two-Pore-Channel 2 (TPC2), stimulated by NAADP agonist but not by phosphoinositide PI(3,5,)P2 agonist. These treatments neutralize lysosomal pH. Moreover, luminal pH changes are also accompanied by changes in ferrous iron availability, though classical ferroptosis inhibitors do not prevent a delayed hair cell death. These findings reveal that lysosomal-driven delayed hair cell death is ferroptosis independent, suggesting that toxicity relies on a distinct mechanism that based on the internal conditions of the lysosomal compartment.

## INTRODUCTION

Mechanosensory hair cells in the inner ear are complex and highly specialized cells that fulfill a pivotal role in hearing and balance ^1^. The importance of these cells is evident by the large number of mutations affecting their development and function that render partial or total hearing loss in humans ^2,3^. Hair cells are also susceptible to damage from certain therapeutic agents, like the aminoglycoside (AG) family of antibiotics^4–6^. Even with these known side effects, AGs are still considered a standard-of-care for many hard to treat diseases like Pseudomonas infections associated with cystic fibrosis ^7–10^, multidrug-resistant tuberculosis and non-tuberculous mycobacterium ^11–14^. The incidence of hearing loss in patients undergoing these treatments can vary from 20-63%. Vestibular issues are also described, though likely is underreported due to absence of widespread clinical methods to determine vestibular function thresholds ^14^. Even though clinical use of AGs in the US has decreased over time, the side effects from these drugs are still a public health concern as AG ototoxicity is estimated to be the most prevalent cause of preventable global hearing loss, generating an estimate of 19.6 million new cases each year ^15^. While AG toxicity is not limited to hair cells, ototoxicity is permanent in humans as these cells are unable to recover from damage ^16^. Thus, understanding the mechanisms by which they damage hair cells may reveal potential therapeutic avenues to make these drugs safer.

The zebrafish has emerged as a valuable model to study mechanosensory hair cells. Zebrafish hair cells closely resemble mammalian hair cells at a molecular level and, due to their transparency, accessibility, and genetic tools available, have become a robust model to study deafness and vestibular dysfunction in vivo ^17,18^. Hair cells in the zebrafish lateral line are grouped in clusters of cells called NMs located on the surface of the body, and are an ideal model to study hair cell ototoxicity due to their easy accessibility and visualization. These hair cells have been shown to be highly susceptible to ototoxic therapeutic drugs, including AGs and cisplatin ^5,19–21^. Moreover, the zebrafish lateral line has been shown useful not only to screen for ototoxic drugs, but also that offer protection against damage ^22,23^.

Our group and others have shown that different AGs are able to trigger distinct temporal hair cell death responses suggesting distinct ototoxicity mechanisms ^5,21^. Acute hair cell death, caused by exposure to AGs like neomycin occurs rapidly, and is triggered by Ca^2+^ release from the endoplasmic reticulum into the mitochondria ^19,24^. This calcium imbalance causes a loss in mitochondrial integrity and an increase in mitochondrial reactive oxygen species (ROS) production resulting in hair cell death ^19,25^. Treatment with the mitochondrial-targeted antioxidant, mitoTEMPO, confers protections against this type of death. Other ototoxic AGs, like gentamicin or the closely related AG G418, result in a delayed cell death after exposure. The ototoxic effects of these AGs are independent of mitochondrial-related pathways ^21^. These AGs are rapidly compartmentalized into vesicles and accumulate in lysosomes ^20,21,26–28^. Treatment with lysosomotropic agents like glycyl-L-phenylalanine 2-naphthylamide (GPN) confers protection against delayed cell death, but not acute death (Wu *et al.*, 2024). These studies suggest that there are separate intracellular loci for each death mechanism: the mitochondria for acute death and the lysosome for delayed death.

In the present study we explore further the mechanisms underlying lysosomal regulation of hair cell death. We aimed to study the vesicular pathways that compartmentalize AGs in hair cells, as well as mechanisms by which lysosomes influence delayed cell death. Specifically, we show that, in zebrafish hair cells in vivo, autophagocytosis is the main compartmentalization mechanism by which AGs enter into lysosomes. We find that activation of lysosomal Two Pore Channel 2 (TPC2) through the NAADP agonist, TPC2-A1-N alters hair cell susceptibility to AGs. This protective effect is only NAADP dependent, as activation of TPC2 through a PI(3,5)P_2_ pathway doesn’t change the survival outcome. Lysosomal modifying drugs induce neutralization of endolysosomal pH, as well as reduction in available ferrous iron in cells. Inhibition of ferroptosis didn’t results in changes in susceptibility during a delayed cell death, but it did protect against an acute cell death. Our work further ratifies the distinct and complex mechanisms influencing AG ototoxicity, and shines light into the compartment-specific pathways influencing cell death.

## MATERIALS AND METHODS

### Animals

Experiments were conducted on 5–7 days post fertilization (dpf) larval zebrafish. Larvae were raised in embryo medium (EM; 14.97mM NaCl, 500μM KCL, 42μM Na_2_HPO_4_, 150μM KH_2_PO_4_, 1mM CaCl_2_ dihydrate, 1mM MgSO_4_, 0.714mM NaHCO_3_, pH 7.2) at 28.5°C. All wildtype animals were of the AB strain. Zebrafish experiments and husbandry followed standard protocols in accordance with University of Washington Institutional Animal Care and Use Committee guidelines. We used four different transgenic lines in this study. The line *Tg(myosin6b:EGFP-Rab7a)^w272Tg^*was used to label late endosomes and lysosomes in hair cells ^21^. Similarly, to label early endosomes we generated a new transgenic line *Tg(myosin6b:EGFP-Rab5ab)* using the Tol2 transposon ^29^ and the hair cell-specific myosin6b promoter ^30^ to direct expression of EGFP fused 5′ to the zebrafish *rab5ab* gene ^31^. To label autophagosomes we generated a new transgenic line *Tg(myosin6b:mScarlet-LC3)* ^32^. Lastly, using the tools mentioned above we also generated a transgenic line to label late endosomes and lysosomes using *Rab7a* but fused to the mScarlet. In this case we also generated point mutations onto the plasmid to create dominant negative (Rab7a^(T22N)^) and dominant active (Rab7a^(Q67L)^) forms of Rab7a, and generated the respective transgenic lines. Transgenic lines were maintained as single lines and replenish by outcrossing them to WT fish. To obtain double transgenic embryos, adults from each independent line were crossed and larvae were screened for double positive signal on hair cells.

### Dose response curves

For dose response curve experiments, we used two protocols to expose larvae to AGs. All protocols treated 7-10 larvae per well in 48-well plates (Nunc^TM^ Cell-Culture Treated Multidishes48, Thermo Fisher Scientific, 150687). For acute cell death experiments (1h) larvae were exposed to neomycin 0-200µM (Sigma-Aldrich, N1142) for 1h. Neo was then washed twice with 500µL of EM, incubated 1h extra hour in 500µL of EM, and euthanized with 1.3% MESAB (MS-222; ethyl-m-aminobenzoate methanesulfonate; Sigma-Aldrich, A5040) before fixation (see below) for hair cell analysis. For delayed cell death experiments (1+23h) larvae were exposed to gentamicin (Sigma-Aldrich, G1397) or G418 (Sigma-Aldrich, A1720) 0-200 µM for 1h. AGs were then washed twice with 500µL of EM followed by an incubation with 700µL of EM for 23h, euthanized and fixed for hair cell analysis.

Instances where ferroptosis inhibitor drugs are tested, larvae were pre-incubated for 2h in 300µL of 20µM Ferrostatin-1 (Cayman Chemicals, 17729), or the respective volume of dimethyl sulfoxide (DMSO; Sigma-Aldrich, D2653). Then neomycin or G418 was added to the well for 1h. Larvae were washed once with 300µL media containing Ferrostatin-1, and incubated for 1h or 23h in 700µL of Ferrostatin-1 or DMSO. Larvae were then euthanized and fixed for hair cell analysis. Similarly, to test if lysosomal proteases can drive delayed cell death, larvae were incubated with the same protocol just described using a mix of E64d (Aloxistatin; MedChemExpress, HY-100229) and Pepstatin A (Sigma Aldrich, P5318) at a concentration of 50µM each, and the respective DMSO volume for the control condition. Larvae were then euthanized and fixed for hair cell analysis.

To test the TPC2 agonist TPC2-A1-N (Sigma-Aldrich, SML3562) and TPC2-A1-P (MedChemExpress, SML3700) compounds, larvae were preincubated for 1h in 3µM TPC2-A1-N and 10µM TPC2-A1-P, respectively. These drugs were then washed once with 500µL of EM, and replaced with 300µL of EM where AGs were then added for 1h. After AG exposure wells were washed twice with 500µL with EM, and incubated in 700µL EM for 1h or 23h respectively, then euthanized and fixed for hair cell analysis.

To inhibit the effects of TPC2-A1-N a modified dose response curve protocol was designed. In short, each group of fish was treated for 1h windows per each drug group with the aim to treat all groups with AG during the same window of time. For the group containing trans-Ned-19 and TPC2-A1-N, larvae were incubated in 300µL of trans-Ned-19 (Tocris, 3954) at different concentrations for 1h. Then, this was replaced with 300µL of a new solution containing trans-Ned-19 with 3µM TPC2-A1-N for another 1h. This solution was then washed out twice with 500µL with EM, and G418 (50µM final concentration) was added to each well for 1h. Lastly, this solution was then washed out twice with 500µL with EM, and larvae were incubated for 23h on 700µL of EM before being euthanized and fixed for hair cell analysis. In the case of the DMSO and trans-Ned-19 control groups, each respective solution was added as 300µL for 1h, then washed out twice with 500µL with EM, and G418 was added to each well for 1h. This solution was then washed out twice with 500µL with EM, and larvae were incubated for 23h on 700µL of EM before being euthanized and fixed for hair cell analysis. EM control group was incubated, washed, and maintained EM solution for the complete experiment using similar volumes as described before.

### Hair cell analysis

After completing euthanization, larvae were fixed with 4% paraformaldehyde (PFA; Thermo Fisher, A16163) in 0.1M Phophate-buffered saline (PBS; pH 7.4) over night in a slow-moving rocker. Larvae were then washed 3x with PBS for 15min at RT and blocked in PBS supplemented with 0.1% Triton-X 100 (PBS-T) and 5% normal goat serum (Sigma-Aldrich, 11H280) for 1–2 h at RT. Samples were incubated in mouse anti-parvalbumin antibody (1:400; EMD Millipore, MAB1572) in blocking solution (PBS-T and 1% normal goat serum) at 4°C overnight. Then samples were rinsed with 3x with PBS-T for 15min at RT, and subsequently incubated in goat anti-mouse antibody conjugated to Alexa 488 (1:500; Thermo Fisher, A-11001) in blocking solution (PBS-T and 1% normal goat serum) at 4°C overnight or 4h at room temperature (RT). Larvae were rinsed with PBS-T 3x for 15min at RT, then with PBS 3x for 15min at RT, and mounted between coverslips with Fluoromount G (SouthernBiotech, 0100-01). Coverslips were held apart by 4 drops of nail polish. Hair cells were counted using a Zeiss Axioplan 2 epifluorescence microscope with a Plan-NEOFLUAR 40x/0.75 NA objective (Zeiss). Counts were performed on four NMs per fish [SO1, SO2, O1, OC1^33^] and summed to arrive at one value per fish. Hair cell counts were performed for 7–10 fish per treatment condition.

### AG and Dextran uptake

To study endocytic activity in hair cells we decided to use dextran polysaccharides (10kDa) linked to an Alexa Fluor 647 moiety as a tracer (Thermo-Fisher, D22914). 10 larvae at a 5dpf stage were transferred to individual wells on a 24-well plates (Falcon, 351147). All EM was taken out and replaced with 800µL of a 400µg/mL of a dextran working solution dissolved in EM. Larvae were incubated in this solution for 5min, or 15min with 15min in EM washout. Then, with the minimum amount of solution, larvae were then transferred to a new 60x15mm petri dish (Falcon, 351007) containing 15mL EM+0.02% MESAB for 5min for anesthetize. The remaining dextran working solution was then transferred onto the next experimental group. Groups of 3 larvae were imaged (described below) using a glass-bottom imaging plate containing 1.5mL of EM+0.02% MESAB. 3 NM per larvae, from a total of 5-7 larvae per condition were imaged.

In order to assess AG accumulation and pH changes in hair cell vesicles we conjugated G418 to BODIPY 650/665-X NHS Ester (Succinimidyl ester; ThermoFisher, D10001) as well to pHrodo Green STP Ester (Thermo Fisher, P35369) following protocols for gentamicin labeling ^34,35^ with previously described modifications ^36^. 10 5dpf larvae per condition were transferred to individual wells on a 24-well plates (Falcon, 351147). For experiments involving TPC2-A1-N and TPC2-A1-P, larvae were preincubated in 1mL of solution at 3µM, 10µM, respectively. Drug solutions were then wash once with 500µL of EM, and larvae were incubated then in 2500µL of EM with 50µM G418. Larvae were then transferred to a new 60x15mm petri dish (Falcon, 351007) containing 15mL EM+0.02% MESAB for 5min for anesthetize. Larvae were imaged (described below) using a glass-bottom imaging chamber containing 1.5mL of EM+0.02% MESAB. For experiments involving glycyl-lphenylalanine 2-naphthylamide (GPN; Cayman, 14634), larvae were co-incubated in 2500µL of EM containing G418 and 250µM GPN for 1h. Larvae were then transferred to a new 60x15mm petri dish (Falcon, 351007) containing 15mL EM+0.02% MESAB for 5min for anesthetize and imaged. Bafilomycin A1 has been reported to affect uptake of AGs, so larvae were exposed to 1h of G418, then 500µL of EM, to finally incubated 1000µL of 100nM Bafilomycin A1 (Baf. A1) for 1h ^21^. For experiments involving GPN and Baf. A1, larvae were imaged with these drugs still in the media. For all experiments measuring pH within vesicles, a 1:1 mix of G418-Bodipy and G418-pHrodo green were prepared before adding it to media containing the samples for 1h. For all conditions 3 NM per larvae, from a total of 5-7 larvae were imaged.

### Hair cell iron levels

To measure ferrous iron level in hair cells 10 5dpf WT larvae per condition were transferred to individual wells on a 24-well plates (Falcon, 351147). Larvae were incubated in 1mL of 3µM TPC2-A1-N or 250µM GPN^37^ for 1h. Then larvae were washed once with 500µL of EM, and replaced with 400µL of EM with 100µM Lyso-FerroRed (Dojindo, L270) for 30min. Larvae were then transferred to a new 60x15mm petri dish (Falcon, 351007) containing 15mL EM+0.02% MESAB for 5min for anesthetize. Larvae were imaged (described below) using a glass-bottom imaging chamber containing 1.5mL of EM+0.02% MESAB. For all conditions 3 NM per larvae, from a total of 5-7 larvae were imaged.

### Imaging

To image individual NM, groups 3 larvae were transferred to a glass-bottom chamber (Ibidi, m-Slide 2) containing 1.5mL of EM+0.02% MESAB, and hold in place using a small piece of 1 x 1 mm net, and two slice tissue harps one on top of the other (Warner Instruments, SHD-26GH/10 and SHD27LH/10). NM were imaged using a Zeiss LSM 980 Airyscan 2 inverted microscope, using a water-immersion LD LCI Plan-Apochromat 40x/1.2NA Imm autocorr DIC objective (Zeiss). Automated correction collar was set to a relative position of 75% for all images. 6 or 7x digital zoom was used depending on the experiment. Z-stack span was defined as 110-120 slices, keeping an interval of 0.21μm for optimal sampling, confirmed by Zen Blue 3.7. Images were obtained using a line-scan AiryFAST SR-4Y scanning method. Sensor gain values, as well as laser intensity, were defined for each experiment, but always trying to minimize photobleaching, reduce unwanted noise, and respecting the optimal values recommended by the manufacturer. These settings were kept identical between technical replicates. The laser line was aligned to the Airyscan sensor array every time a glass-bottom chamber was switch.

### Image Processing

For colocalization analysis 3D Airyscan processing of all images was performed on Zen Blue 3.7, with a strength value of 4.5 for all channels. These images were then run through a FIJI ^38^ macros to obtain Pearson’s correlation values. In short, czi formatted multichannel z-stacks were imported through BioFormats importer plugin ^39^, then background was substracted on each slice using a rolling ball method with a size of 50. Using the GFP-signal channel, a 3D mask was created by applying a gaussian filter of sigma value 2, and then thresholded using a minimum error method. This mask was post-processed to fill holes within. Colocalization was obtained through the FIJI plugin Coloc 2, including the previously generated mask, a bisection method for threshold regression, and other values as default. Pearson’s correlation values (above threshold) were then taken for further statistical analysis. To analyze Lyso-FerroRed intensity in hair cells, czi formatted multichannel z-stacks were imported through BioFormats importer plugin. Maximum intensity projections were created, regions of interest were drawn manually. Background was subtracted and integrated density was calculated.

### Automated segmentation

To measure AG accumulation and changes in pH within vesicles we used a similar protocol as described in with slight updates. Using a FIJI macros, czi files were converted to TIFF. Then, images were imported and vesicles were segmented using the Allen Cell and Structure Segmenter ^40^. Intensity normalization function was applied to both channels, a gaussian filter of sigma value 1 is applied to the z-stack, and are these new images that are subsequently used for segmentation. Vesicles were segmented using the 2D slice-by-slice dot wrapper and 2D filament wrapper functions from the Allen Cell Classic Segmenter package with the following parameters: for the G418-Bodipy channel (Figure S3), Dot segmenter parameters: scale_spot_1 = 4, cutoff_spot_1 = 0.12, scale_spot_2 = 2, cutoff_spot_2 = 0.1, and Filament segmenter parameters: scale_fil_1 = 3, cutoff_fil_1 = 1, scale_fil_2 = 1, cutoff_fil_2 = 0.9. The two segmenter results were fused together to create a single vesicle mask, and small objects (bellow a 60 pixels of size) were removed. To avoid some segmentation artifacts, holes within it were filled with a maximum size of 250 pixels. Watershed segmentation was then applied to create individual masks out of the vesicle network, and vesicles were labeled using the label function from scikit-image ^41^. This method allows for good an overall individual vesicle mask generation, though we acknowledge that vesicular tubule structures become over-segmented in these conditions. Unlike our previous method, we are unable to segment the hair cells within the NM, though using a far red signal we notice very little non-specific segmentation outside of it. A background mask was created by inverting the vesicular mask. Using these masks, vesicle and background properties were calculated using regionprops_table function from scikit-image. Properties for single vesicles and background were concatenated into a single table and exported to a Microsoft excel file. Vesicle number, volume (voxel number), and intensity data, within these files was organized and background corrected. Fluorescent intensity grey values were normalized to vesicle area. These integrated density values were then averaged per each NM and used to calculate the average integrated density ratio per NM.

### Statistics

Mann-Whitney tests, T-tests, one way or two-way ANOVA with post-hoc testing was performed using GraphPad Prism 11. For graphical presentation of dose response curves, data were normalized to untreated controls such that 100% represents hair cell survival in control animals. Results were considered significant if p ≤ 0.05 with levels of statistical significance shown in figures as follows: ns; P > 0.05, ∗; P <= 0.05, ∗∗; P <= 0.01, ∗∗∗; P<= 0.001, ∗∗∗∗; P<0.0001.

## RESULTS

### The autophagocytic pathway as the main AG compartmentalization route in zebrafish hair cells

Accumulation of AGs in lysosomes has been proposed to occur through two distinct mechanisms: endocytosis and lysosomal trafficking^20,42,43^ or through autophagy after channel entry^20,44,45^. To follow AG trafficking through the endocytic pathway, we labeled late endosomes/lysosomes with Tg(myo6b:EGFP-Rab7a) ^21^, as well as create a new transgenic line, Tg(myo6b:EGFP-Rab5ab), to label early endosomes ^46,47^. Vesicles observed in the new Rab5ab are in line with what others have observed in the past, with Rab5ab vesicles being enlarged and located apically in the cell ^48^.

To assess endocytosis in zebrafish hair cells, we exposed larvae to dextran (10kDa) linked to an Alexa 647 fluorophore ^49^. After 5min pulse of dextran, we observed some accumulation inside Rab5ab-positive vesicles inside hair cells, but little to no accumulation in Rab7a-positive vesicles [Fig. 1A]. After extending uptake time to 30min, we saw a stronger Rab5ab accumulation, with a larger population of Rab5ab vesicles containing dextran [Fig. 1B]. We observed increased accumulation in Rab7a-positive vesicles, consistent with maturation of the endocytic compartment [Fig. 1B]. We note that our images also show significant uptake of dextran by the GFP-negative support cell population around hair cells (MIP column on Fig. 2A,B) suggesting, different levels of endocytic activity among these two cell populations.

**Figure 1:**
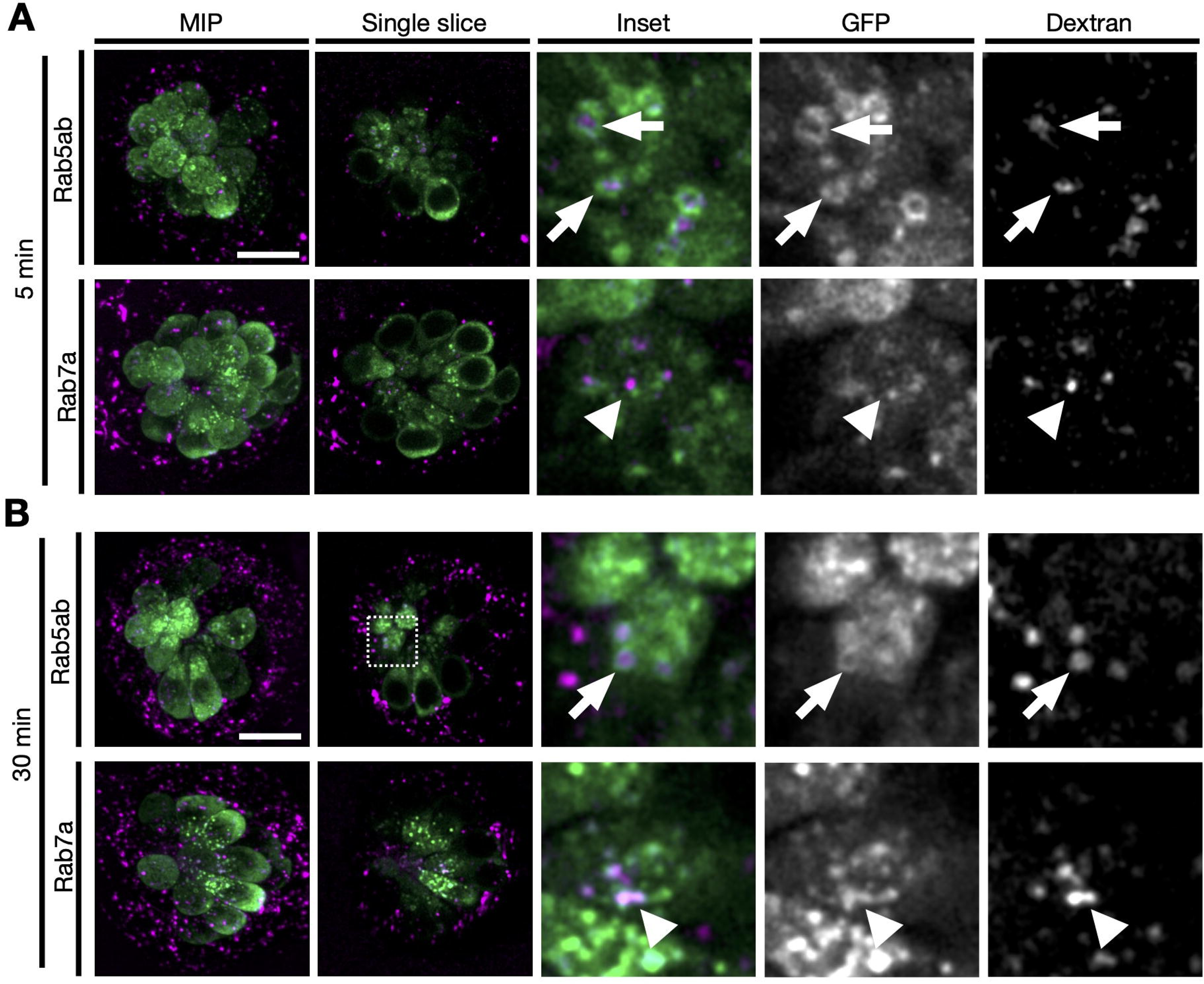
Endocytosis is the main dextran internalization pathway. **A)** 5dfp transgenic larvae, expressing EGFP-Rab5ab or EGFP-Rab7a (green) in hair cells, were pulsed with 400µM Dextran-Alexa647 (magenta) for 5min before imaging. Maximum intensity projections (MIP) of the whole NM are shown. Representative single slice images are selected to show Rab5ab/Rab7a-positive and dextran-loaded vesicles inside hair cells. Many Rab5ab-positive vesicles show some accumulation of dextran during early uptake (arrow). Rab7a-positive vesicles, appear to be loaded or associated with dextran-containing vesicles (arrowhead). **B)** Larvae were exposed for 30min to dextran before imaging. Longer exposure shows clear accumulation of dextran in Rab5ab-positive vesicles (arrow), as well as in Rab7a-positive vesicles (arrowhead). Scale bar: 10μm.

**Figure 2:**
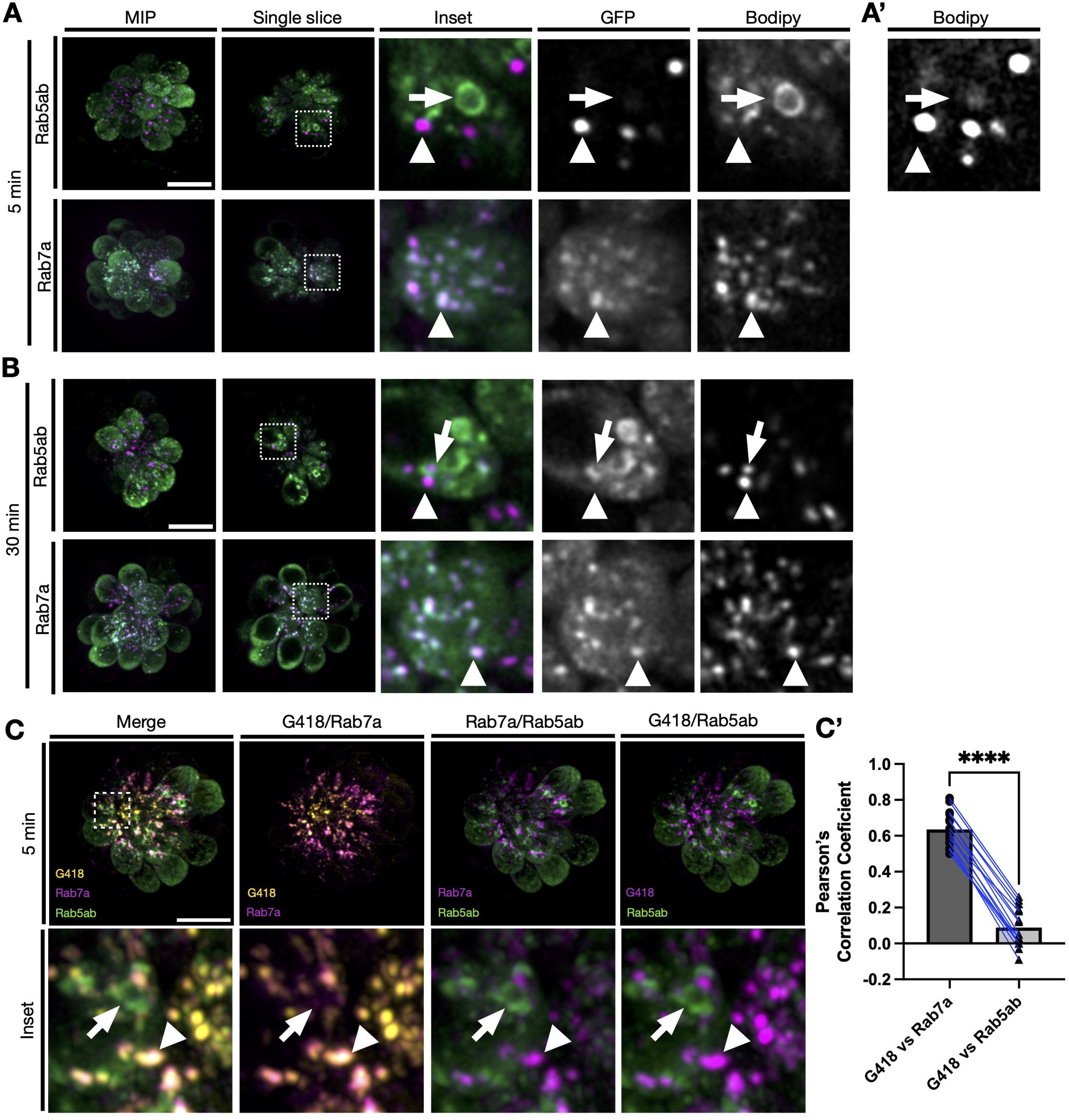
Early uptake of G418 accumulates in late endosomes/lysosomes. **A)** 5dfp transgenic larvae, expressing EGFP-Rab5ab in hair cells, were pulsed with 50µM G418-Bodipy (magenta) for 5 minutes before imaging. Maximum intensity projections (MIP) of the whole NM are shown. Representative single slice images show little accumulation Rab5ab-positive vesicles. Rab7a-positive vesicles (arrow) show high accumulation of G418 in them (arrowhead). **A’)** A few Rab5ab vesicles show dim G418-Bodipy fluorescence, visible only if histogram levels are increased. **B)** Same transgenic lines were pulsed for 15min to G418-Bodipy, washed, and incubated for 15min in EM before imaging. Longer exposures show some accumulation of G418 in Rab5ab-positive vesicles (arrow). In a similar way as before, most accumulation happens inside Rab7a-positive vesicles (arrowhead). **C)** 5dfp double transgenic larvae, expressing EGFP-Rab5ab (green) or mScarlet-Rab7a (magenta) in hair cells, were pulsed with 50µM G418-Bodipy (yellow) for 5min before imaging (Merge). Inset images show that G418 colocalize almost exclusively with Rab7a (arrowhead), when compared to Rab5ab (arrow). **C’)** Pearson’s correlation coefficient show a strong colocalization between G418 and Rab7a signal. Rab5ab and G418 signal shows little to no correlation. Stats: Mann-Whitney t-test. Scale bar: 10µm.

We next tracked the endocytic accumulation of G418 linked to a Bodipy650/665-X fluorescent moiety over time (G418-Bodipy). After a 5-minute exposure, we saw bright vesicles loaded with G418. Almost all of these vesicles were Rab5ab-negative [Fig. 2A], although a few Rab5ab-positive vesicles containing small amounts of the AG were observed [Fig. 2A’]. By contrast, almost all G418+ vesicles were labeled with EGFP-Rab7a, indicating that the AG was accumulating in late endosomes and lysosomes [Fig. 2A]. By extending uptake to 15min of G418 exposure followed by an additional 15min incubation time in embryo media (EM), only a small fraction of Rab5ab-positive vesicles are loaded with AGs [Fig. 2B]. As previously, nearly all G418+ vesicles were Rab7a-positive [Fig. 2B]. Moreover, 3D colocalization analysis between G418 and Rab7a/Rab5ab-positive vesicles confirms very little accumulation of G418 in early endosomes during early phases of uptake [Fig. 2C, C’]. This data suggests that G418 is being compartmentalized by other vesicular pathways.

Previous studies have demonstrated that autophagy can play a role during AG uptake and toxicity ^44,45,50^. We hypothesize that, in our model, this pathway could close the gap between entry of AGs through the MET channel and their accumulation in late endosomes/lysosomes. In order to investigate this, we generated a zebrafish transgenic line, *Tg(myo6b:mScarlet-LC3),* to label autophagosomes in hair cells ^50,51^. Similarly to hair cells expressing EGFP-Rab7a, this fluorescent probe labels compact, individual vesicles, most of them on the apical side of cells. Exposure of this transgenic line to G418 shows high accumulation in this vesicular compartment [Fig. S1A]. Double expression of mScarlet-LC3 and EGFP-Rab7a in hair cells show that, as expected, these compartments overlap, as many vesicles are double labeled with both probes. A small subset of Rab7a vesicles do not co-label with LC3, suggesting they may be from the endocytic pathway instead of formed through autophagy [Fig. 3A]. When exposed to G418, these three signals - LC3, Rab7a, and G418 - colocalize within hair cells [Fig. 3B]. These data suggest that, for drugs like G418 that trigger a delayed hair cell death, autophagocytosis is the main route towards the mature vesicular compartment.

**Figure 3:**
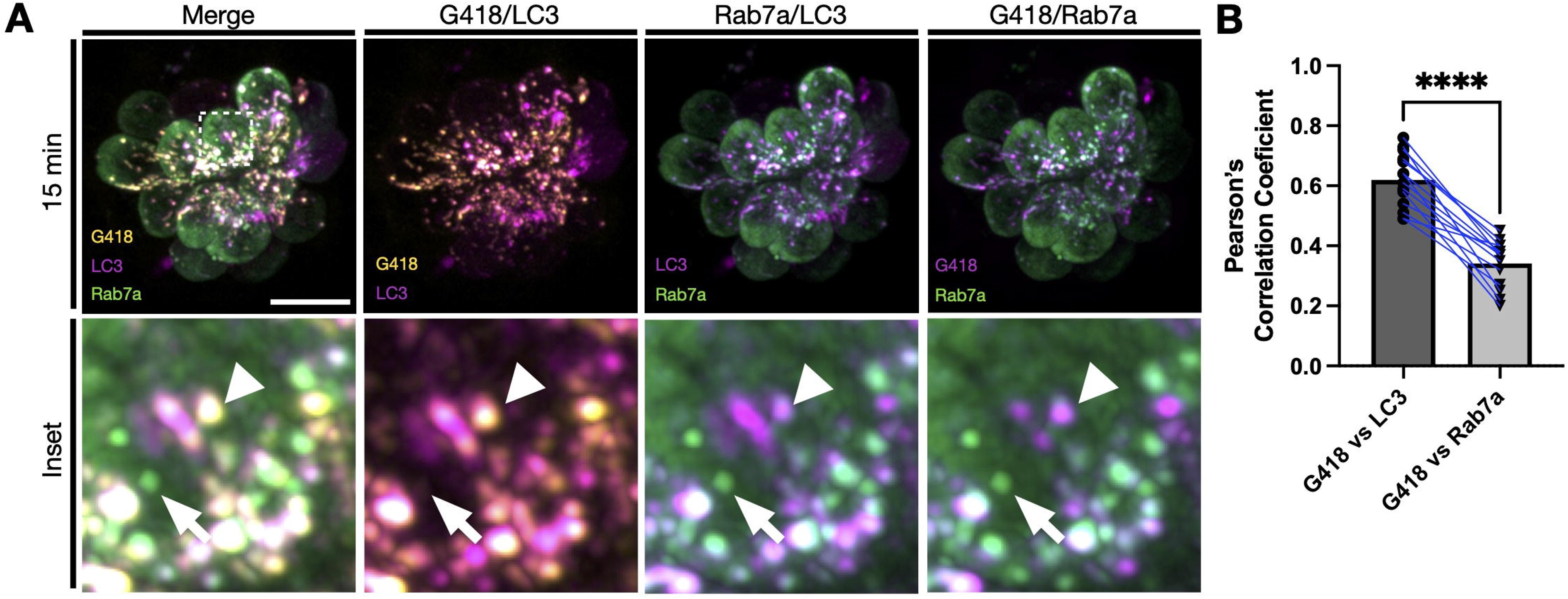
Autophagocytosis as G418 capture and accumulation pathway. **A)** 5dfp double transgenic larvae, expressing mScarlet-LC3 (magenta) and EGFP-Rab7a (green) in hair cells, were pulsed with 50µM G418-Bodipy (yellow) for 15 minutes before imaging (Merge). Individual dual-channel images are pseudocolored to show high accumulation of G418 in LC3-positive vesicles (arrowhead). Rab7a-positive vesicles show accumulation of G418, though some show no colocalization with other signals (LC3 or G418; arrow). **B)** Pearson’s correlation coefficient show a strong correlation between LC3 and G418 signal, and a slightly lower for Rab7a and G418 signal. Stats: Nonparametric Mann-Whitney t-test. Scale bar: 10μm.

Rab7 has been extensively described to be a mayor player during early-to-late endosomal maturation, lysosome acidification, transport, as well as autophagosome/lysosome fusion ^46^. We hypothesized that interfering with Rab7a function would change hair cell susceptibility to AG and accumulation in lysosomes. With this in mind, we created two new transgenic lines expressing dominant negative (Rab7a^T22N^), or dominant active (Rab7a^Q67L^) mutations ^31^ in hair cells. Expression of Rab7a^T22N^ results in marked disruption of the vesicle network, as vesicles are found in lower numbers, morphologically rounder and located throughout the hair cell cytoplasm instead of enriched at the apical end of the cell. Expression of Rab7a^Q67L^ results in comparable phenotype as for Rab7a, with slightly higher fragmentation of vesicles, that nevertheless maintain their apical localization [Fig. S1B]. These phenotypes are in line with severally previously described ^46,52,53^. We next tested whether expression of these mutant proteins resulted in dose dependent changes in hair cell death with exposure to G418. For both lines there was no change in the dose response function [Fig. S1C]. These results suggest that Rab7a regulation of lysosomal maturation does not influence AG toxicity in hair cells. Alternatively, due to the reactivity characteristics of AGs, we hypothesized that they could induce Lysosomal membrane permeabilization (LMP) after G418 uptake can result in transfer of cathepsin proteases to the cytoplasm, resulting in necrotic cell death ^54,55^. Inhibition of lysosomal cathepsins by incubation with the protease inhibitors PepstatinA and E64d, shows no protection after exposure to G418 [Fig. 4], suggesting that G418 ototoxicity is not linked to protease activity and release.

**Figure 4:**
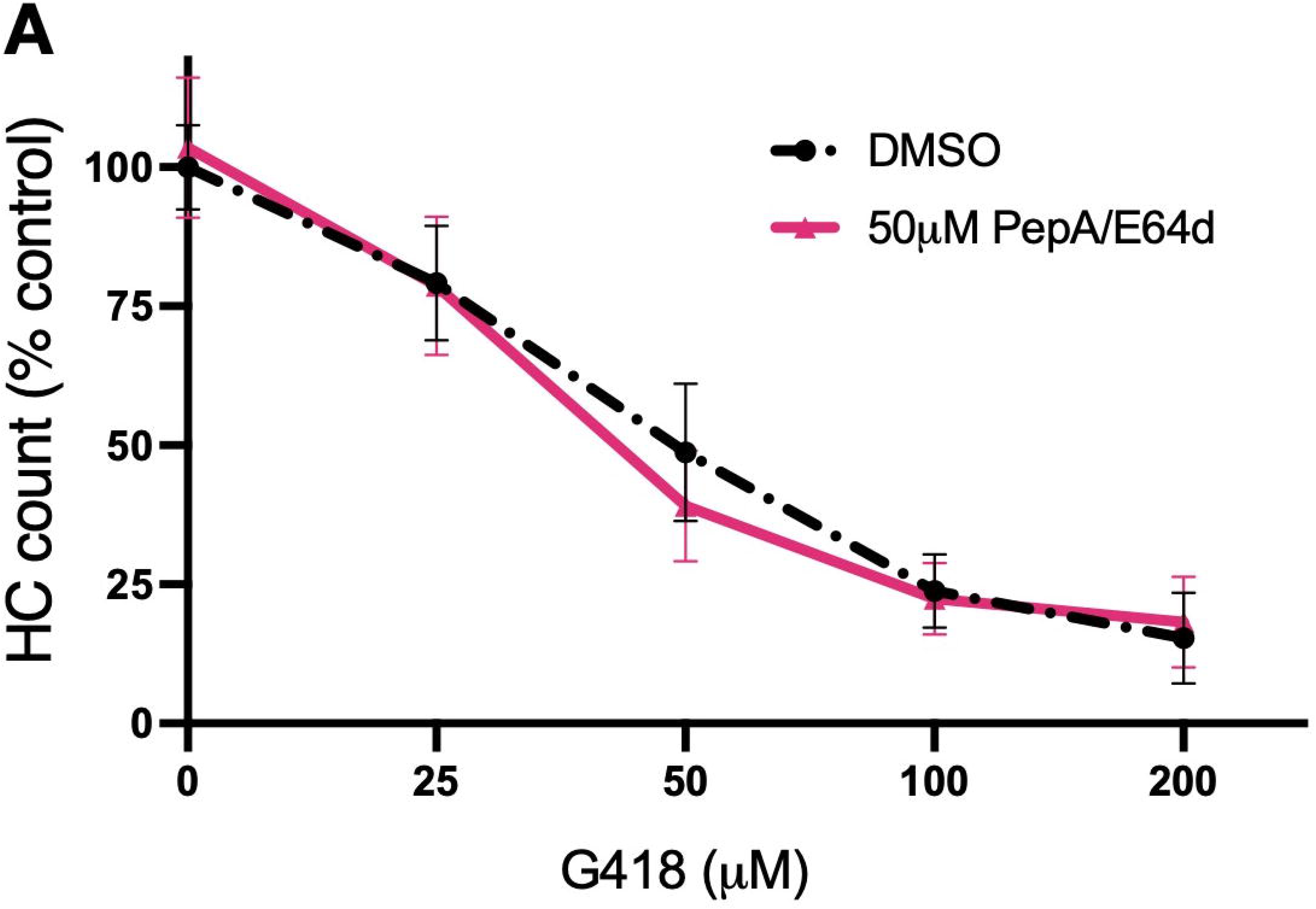
Protease inhibitors don’t protect against delayed hair cell death. **A)** 5dpf WT larvae were co-incubated with 50µM of Pepstatin A/E64d and G418 for 1h at different concentrations. After G418 exposure, larvae were washed and incubated back in PepA/E64d solution for 23h. No differences were found between groups. Stats: Two-way ANOVA, column factor P-value: 0.6280.

### Activation of TPC2 channels protects against delayed hair cell death

We previously demonstrated that interference with lysosomes by exposure to GPN protects hair cells against delayed, but not acute cell death ^21^. GPN has been described to cause multiple alterations to lysosomal homeostasis, perhaps most prominently causing Lysosomal Membrane Permeabilization (LMP) and subsequent mobilization of Ca^2+^ ions out of this acidic compartment^56–58,56–58^. We hypothesized that lysosomal Ca^2+^release could be linked to changes in hair cell susceptibility to AGs. The Two-Pore Channel 2 protein (TPC2) has been described to be expressed exclusively in lysosomes ^59,60^. TPC2 permeability can be altered by two second messenger molecules, Nicotinic acid adenine dinucleotide phosphate (NAADP) and Phosphatidylinositol 3,5 bisphosphate (PI(3,5)P_2_), that influence the release of Ca^2+^ or Na^+^ ions into the cytoplasm, respectively. Gerndt et al., reported two membrane-permeable small molecule agonists for TPC2, TPC2-A1-N and TPC2-A1-P, that respectively activate the NAADP or PI(3,5)P_2_ pathways and in turn, induces the release lysosomal ions ^60^. We found that pre-treatment with TPC2-A1-N for 1h conferred marked protection against G418 [Fig. 5A] or gentamicin [Fig. S2A] but not to neomycin [Fig. 5A], similar to the action of GPN ^21^.

**Figure 5:**
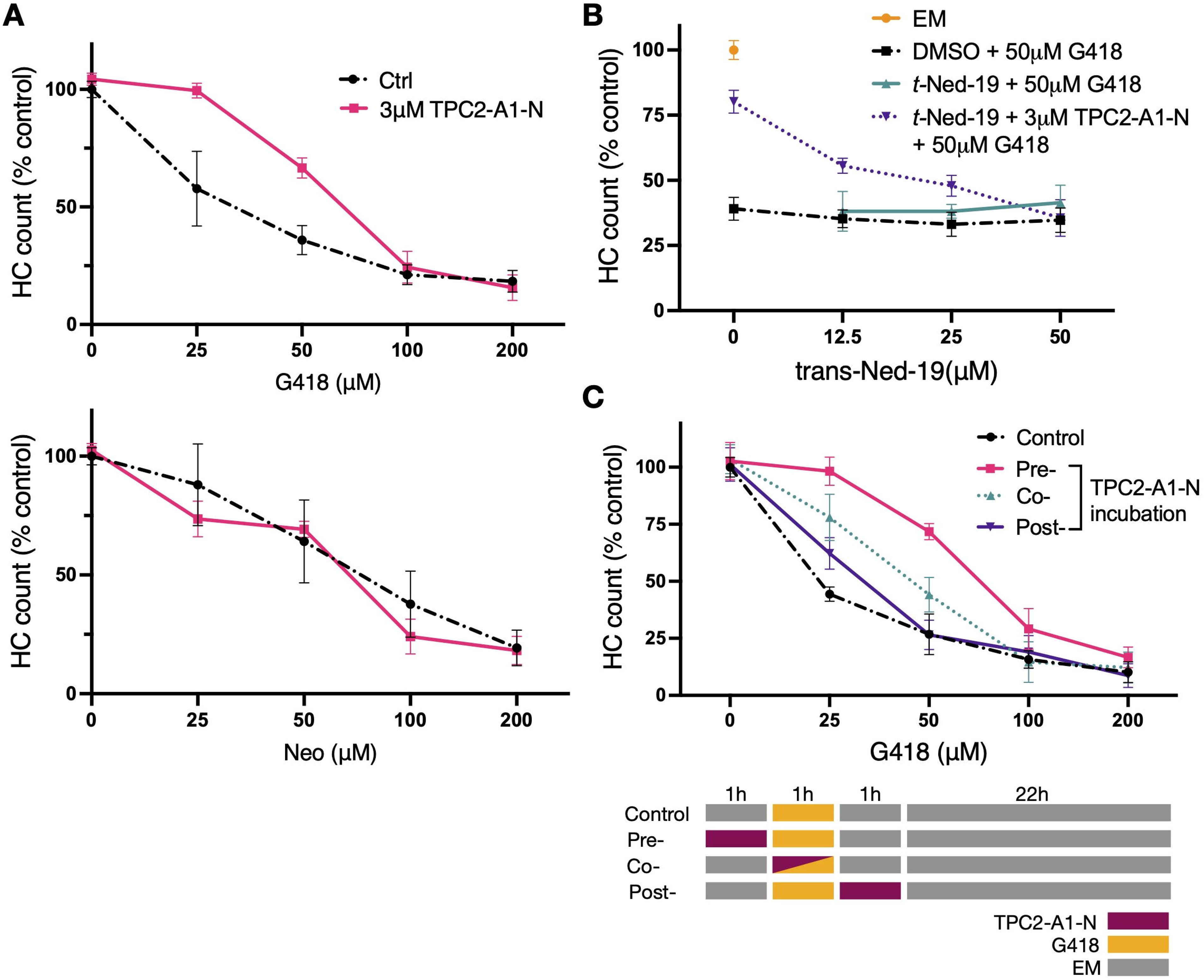
TPC2-A1-N exposure protects against delayed hair cell death. **A)** 5dpf WT larvae were pre-incubated with TPC2-A1-N for 1h, washed out, and incubated for 1h to G418 or neomycin at different concentrations. In the case of G418, after 1h of AG exposure, larvae were washed and incubated in EM for 23h before fixation. For neomycin, after 1h of AG exposure, larvae were washed and incubated in EM for 1h before fixation. Surviving hair cell were counted, and counts were normalized against EM control. TPC2-A1-N protects against G418, but not neomycin. Stats: Two-way ANOVA, column factor P-value: G418: <0.0001; Neo: 0.0671. **B)** 5dpf WT larvae were incubated sequentially with trans-Ned-19 for 1h, 3μM TPC2-A1-N for 1h, washed, and finally 50μM G418 1h. Larvae were then washed with EM and incubated for 23h before quantification. Counts are normalized against EM treated WT larvae. TPC2-A1-N protection is inhibited by trans-Ned-19. No differences were detected between DMSO and t-Ned-19. Stats: Two-way ANOVA, column factor P-value: DMSO vs t-Ned-19: 0.0031 **C)** 5dpf WT larvae were exposed to 3μM TPC2-A1-N at different intervals compared to G418 as shown in the diagram below. Larvae were then washed and incubated in EM for 22h. Surviving hair cell were counted, and counts were normalized against DMSO control. TPC2-A1-N protection is triggered in a time-sensitive. Maximum protection is achieved with pre-incubation, co-incubation conveys reduced protection, and post-incubation shows no differences with control Stats: Two-way ANOVA, Šídák’s multiple comparison test P-value against control: Pre-: <0.0001; Co-: 0.0002; Post-: 0.3719.

By contrast, treatment with TPC2-A1-P offered no protection [Fig. S2B], suggesting that the PI(3,5)P_2_ pathway does not underlie protection. Neither treatment with TPC2-A1-N nor with TPC2-A1-P had an effect on the overall accumulation of AG [Fig. S4A,B]. Unlike GPN ^21^, treatment with either TPC2 drug did not affect endolysosomal vesicle volume or number [Fig. S4C,D]. To further support that activation of TPC2 through the NAADP pathway confers protection, we assessed the effectiveness of TPC2-A1-N in the presence of trans-Ned-19, a selective blocker of NAADP-dependent Ca^2+^ release ^61,62^. While exposure to trans-Ned-19 alone has no effect on the toxicity of G418, increasing doses of trans-Ned-19 effectively eliminates protection conferred by TPC2-A1-N [Fig. 5B]. Overall, these results suggest that activation of TPC2 through the NAADP pathway protects hair cells against a delayed cell death.

We next tested when stimulation of TPC2 channels was most effective compared to G418 exposure. Preincubation with TPC2-A1-N before G418 exposure renders maximum protection compared to co-incubation with TPC2-A1-N and G418. Exposure to TPC2-A1-N after G418 uptake offers no substantial protection [Fig. 5C]. Overall, our results suggest that activation of TPC2 through the NAADP pathway before uptake of G418 is protective against a delayed, but not acute cell death. This protection is specific to calcium release, as sodium release, through the exposure of TPC2-A1-P has no effects.

### TPC2-A1-N induces lysosomal pH alkalinization in hair cells

NAADP-induced Ca^2+^release, through TPC2, has been previously described to induce lysosomal luminal pH changes ^60,63^. To assess the pH inside G418-loaded vesicles, we developed a ratiometric analysis using a mixture of G418-Bodipy and G418 conjugated to pHrodo green, a pH sensitive dye that increases fluorescence with acidification. This labeling system, allowed us to couple it with an automated 3D vesicle segmentation pipeline to measure volume and pH in parallel in a vast variety of vesicle sizes and intensities [Fig. S3A,B]. We tested our system by exposing hair cells to GPN and to Bafilomycin A1 (Baf. A1), as these molecules have been shown to neutralize lysosomal pH, albeit through different mechanisms ^57^. In line with previous observations, our system shows that GPN and Baf. A1 diminish the number of vesicles and at the same time enlarging their average volume [Fig. S3C,E] ^21,64^. More importantly, the luminal pH of vesicles containing G418 is strongly alkalinized [Fig. S3D,F]. Subjected to the same incubation strategy as before [Fig. 5A], we observe that in all conditions vesicles internalized both probes successfully [Fig. 6A, Fig. S4A,B]. In line with previously described in vitro models, TPC2-A1-N reduces the lysosomal pH of vesicles [Fig. 6B]. On the other hand, exposure to TPC2-A1-P left luminal pH largely unaffected [Fig. 6C]. Overall, our data suggest that early ion release and pH changes within late endosome and lysosomes compartment are able to generate conditions that inhibit G418-mediated ototoxicity.

**Figure 6:**
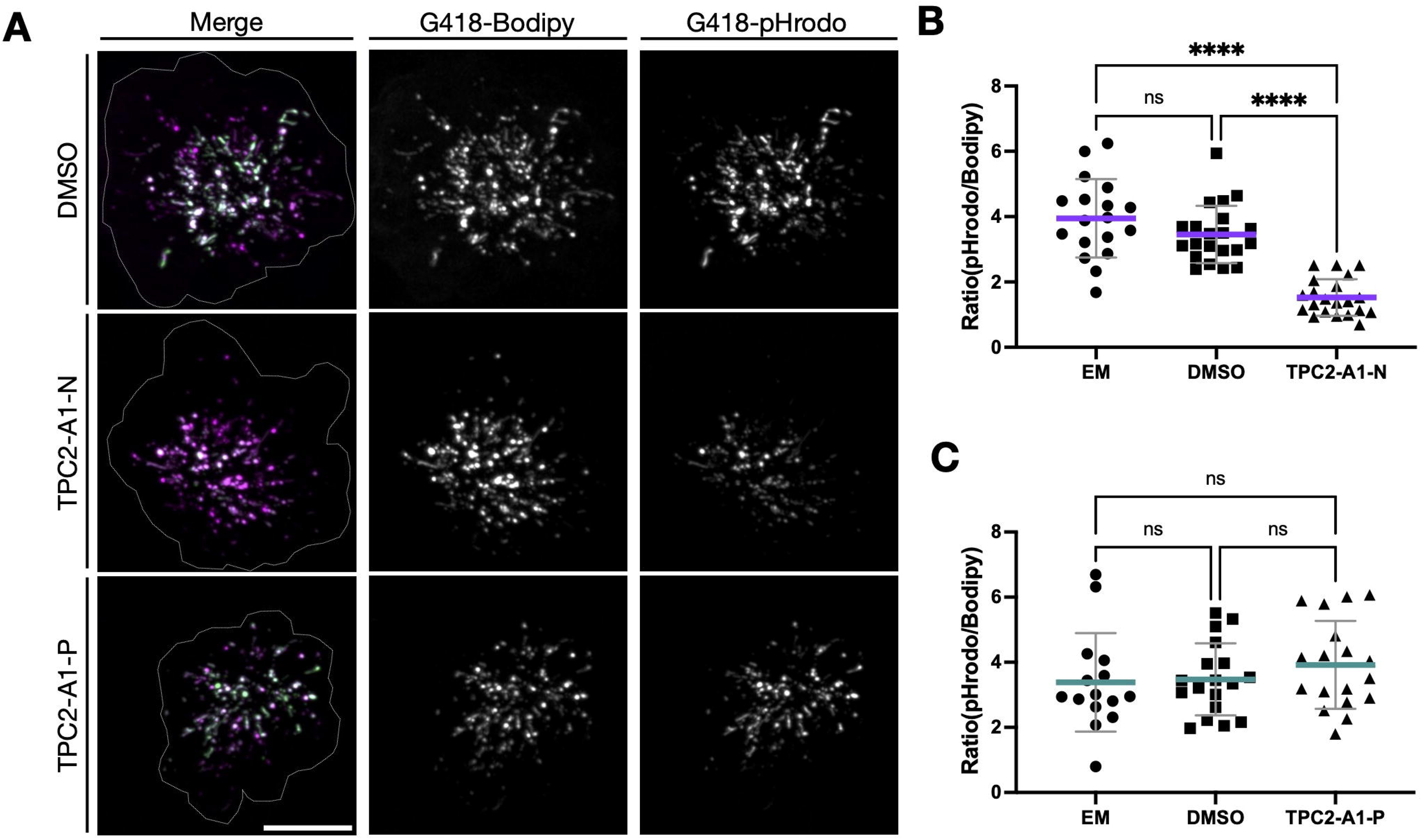
TPC2-A1-N neutralizes pH in vesicular lumen. **A)** WT larvae were incubated with specific agonists for 1h, then washed, and exposed to 50µM of a mixture of equal parts of G418-Bodipy and G418-pHrodo green for 1h before imaging. Representative images show G418-Bodipy (magenta) localizing to a large extent of vesicles in multiple hair cells, whereas high intensity G418-pHrodo green signal localizes in discrete regions of NM (Merge). Only TPC2-A1-N generates a lower pHrodo green fluorescence intensity. Dotted line delineates all hair cells in the NM. Scale bar: 10μm. **B)** Using G418-Bodipy fluorescent signal vesicles were segmented. Normalizing mean intensity by the volume of vesicles, integrated density was calculated on both channels, and mean vesicular ratiometric values were calculated. Each dot corresponds to a NM (average vesicle values), 3 NM per larva. 1h Pre-exposure to TPC2-A1-N neutralizes pH in G418-containing vesicles. Non-param. One-way ANOVA. **C)** Ratiometric values were calculates as previously described. 1h Pre-exposure to TPC2-A1-P doesn’t affect vesicular pH. Stats: Kruskal-Wallis One-way ANOVA, with Dunn’s multiple comparison test.

### Delayed hair cell loss is not altered by drugs that engage lysosomal cell death pathways

Ferroptosis is a form of cell death characterized by accumulation of lipid-peroxides driven by the accumulation of iron ions ^65,66^. Since lysosomes are key regulators of iron homeostasis ^67^ and have been described as a hub for ferroptosis initiation ^68,69^ we tested whether the well-characterized ferroptosis inhibitor Ferrostatin-1 altered the dose-dependent effects of G418. In order to inhibit ferroptosis during all the experimental window, we decided to incubate our samples for 2h, before we expose hair cells to G418 for 1h, and incubate for 23h in clean media containing Ferrostatin-1. We found that exposure Ferrostatin-1 didn’t altered the dose-dependent killing of hair cells by G418 [Fig. 7A]. As described in previous publications, Ferrostatin-1 (and a similar compound, Liproxtatin-1) are able to partially protect hair cells against other ototoxic compounds, like cisplatin or other AGs ^70,71^. Thus, we decided to test if Ferrostatin-1 is able to protect hair cells against an acute cell death triggered by neomycin. In line with previous reports, Ferrostatin-1 is able to induce protection against neomycin, suggesting this compound can protect hair cells against acute cell death, but not delayed one [Fig. 7B]. The lysosomal compartment plays an important role on iron metabolism, and is in here that iron cycles between ferric (Fe^3+^) and ferrous (Fe^2+^) states, needed for multiple cellular processes ^72^. Ferrous iron availability needs to be tightly regulated, because via the Fenton reaction, it can induce damage in protein, nucleic acids and membrane lipids ^54,72^. Changes in pH in the lysosome lumen can alter the balance between ferrous and ferric states, thus affecting ferrous iron availability ^66,73^. We therefore decided to measure the levels of ferrous iron in hair cells after the exposure to drugs that confer protections against G418. Using Lyso-FerroRed, we are able to see that both TPC2-A1-N and GPN, are able to lower the ferrous iron availability [Fig. 7C]. Collectively, these data support a model AGs that trigger a delayed cell death are delivered through autophagy to lysosomes, and it is the internal lysosomal conditions, like pH and iron availability, that influence ototoxicity.

**Figure 7:**
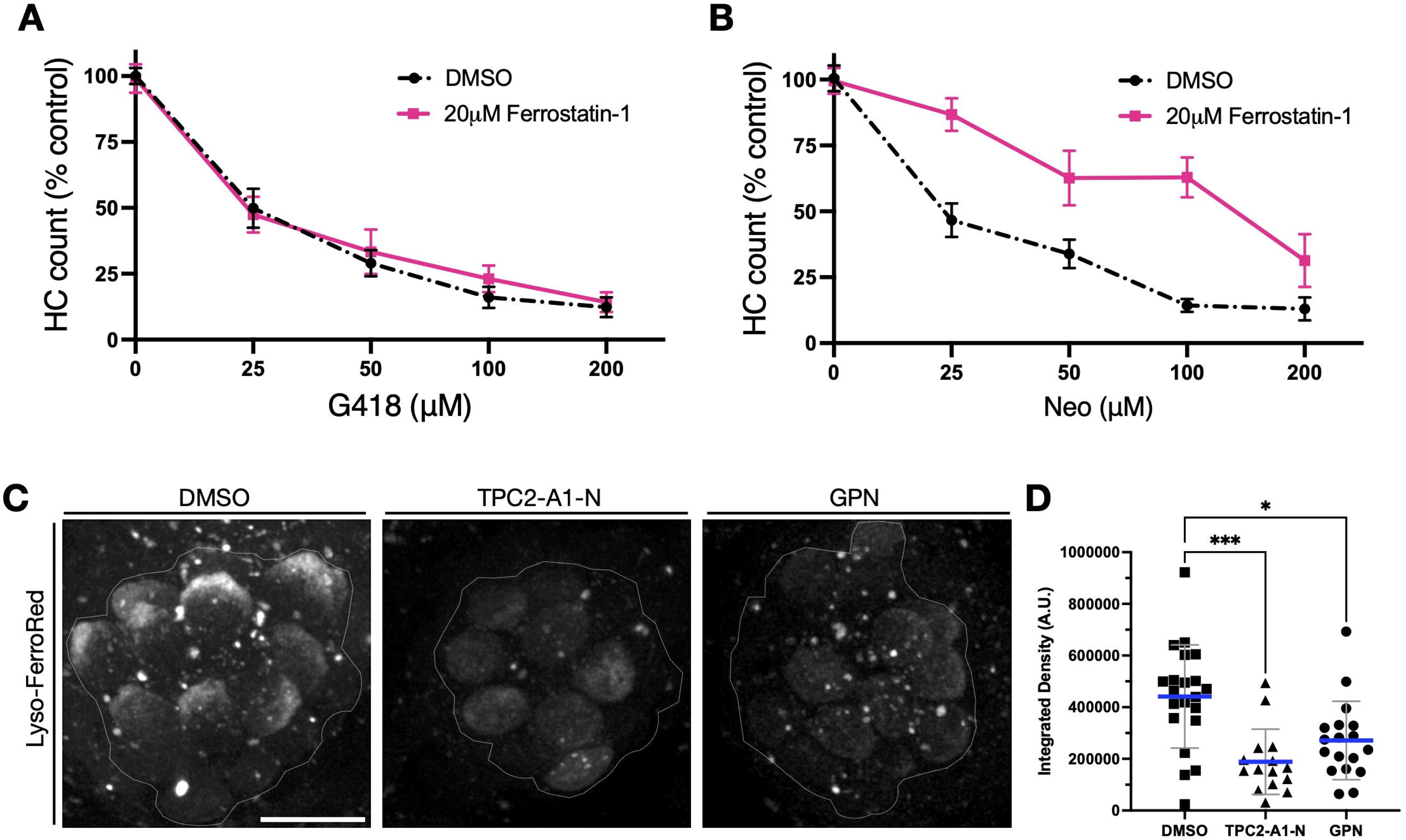
Ferrostatin-1 doesn’t protect against delayed hair cell death. **A)** 5dpf WT larvae were pre-incubated with 20µM ferrostatin-1 for 2h, washout, then incubated 1h with G418 or neomycin at different concentrations. In the case of G418, after 1h of AG exposure, larvae were washed and incubated in EM for 23h. Stats: Two-way ANOVA, column factor P-value: 0.8306 **B)** For neomycin, after 1h of AG exposure, larvae were washed and incubated in EM for 1h. Surviving hair cell were counted, and counts were normalized against DMSO control. Ferrostatin-1 is able to protect against neomycin, but not G418 exposure. Stats: Two-way ANOVA, column factor P-value: <0.0001. **C)** 5dpf WT larvae were incubated 1h with 3μM TPC2-A1-N or 250μM GPN, washed, and incubated with 100μM of Lyso-FerroRed for 30min. The protective agents, TPC2-A1-N and GPN, are able to reduce the concentration of Fe^2+^ in the NM. Dotted line delineates all hair cells in NM. Scale bar: 10μm. **D)** Maximum intensity projections (MIP), were used to manualy draw the hair cell area, and calculate integrated density. TPC2-A1-N and GPN are able to reduce the amount of Fe^2+^ in hair cells. Stats: Kruskal-Wallis One-way ANOVA.

## DISCUSSION

In this study we examine mechanisms underlying a delayed cell death observed in zebrafish lateral line hair cells after AG exposure. We confirm and extend our previous work ^5,20,21^ demonstrating an acute hair cell death during exposure to neomycin and a distinct process that occurs long after exposure to gentamicin or the closely related AG G418. Our studies provide additional evidence that the lysosome is central to delayed cell death, while mitochondria play a critical role for acute cell death ^21,25^.

We provide evidence that autophagy is the major mechanism for accumulation of AGs in lysosomes of zebrafish lateral line. Although apical endocytosis is active in hair cells^20,74,75^, following the trafficking of fluorescent tagged AG in comparison to fluorescent dextran in lateral line hair cells using transgenic markers of the endocytic and autophagic pathways supports autophagy as the major entry of AGs into the vesicular compartment. Work from Zhao and colleagues has implicated autophagy as a central mechanism in AG induced hair cell death in mouse cochlea^44,45,50^, demonstrating that key autophagy proteins GABARAP and MAP1LC3B are necessary for ototoxicity. In these studies, mitochondrial changes and mitophagy were considered the important components downstream of AG exposure, as reduction of the mitochondrial surveillance proteins Pink/Parkin reduced ototoxicity. However, Parkin has been implicated in lysosome-mitochondrial interactions and metabolic homeostasis ^76^, and is also a key regulator of LRRK2, which has central roles in lysosome damage responses ^77^. Future studies will help determine whether Pink/Parkin and LRRK2 regulate hair cell death in zebrafish.

Our initial observations suggest that a canonical ferroptosis pathway does not underlie delayed cell death caused by lysosomal accumulation of AG. We found that treatment with the ferroptosis inhibitor Ferrostatin-1 had no effect on delayed cell death in lateral line hair cells. While Ferrostatin-1 has been described as an inhibitor of the lipoxygenase enzyme complex 15LOX/PEBP1 that results in lysosomal membrane damage^78^, Ferrostatin-1 and the related compound Liproxstatin-1 also have more general antioxidant properties ^79^. Fan and colleagues previously reported that Liproxstatin-1 protected zebrafish hair cells against neomycin exposure ^71^, suggesting it protected against the acute death mechanism. We report here that Ferrostatin-1 protects against neomycin-induced acute hair cell death, consistent with these findings. Our previous studies implicated mitochondrial oxidation underlying acute hair cell death^21,25^. Whether iron-dependent lipid peroxidation occurs in mitochondria as part of this process awaits further study.

Despite our initial findings, we cannot yet rule out that some form of ferroptosis causes the lysosomal-dependent delayed hair cell death that we observe with gentamicin and G418 treatment. Indeed it is becoming increasingly clear that different aspects of ferroptosis vary depending on cell type and condition. Iron-dependent lipid peroxidation is the central mechanism underlying ferroptosis, and in some cases is brought about by inhibition of endogenous control mechanisms that prevent peroxidation. These include glutathione peroxidases GPX4 ^80^, GPX1 ^81^, and the oxidoreductase FSP1 ^82,83^ in different contexts, inhibition of each of these suppressors can trigger ferroptosis. Ferroptosis signals also originate in distinct locations depending on context, including mitochondria, endoplasmic reticulum and plasma membrane (reviewed in Stockwell, 2022^84^). The lysosome has also been implicated as an origin of ferroptosis, with its central role in iron storage and metabolism ^69^. Moreover, ferroptosis can result in heterogeneous forms of cell death ^85^. Work of Schacht and colleagues implicated interactions between iron, gentamicin and phospholipids as a potential mechanism underlying AG ototoxicity ^54,86^, which may occur independently of factors that initiate ferroptosis in other cell types.

Our findings that altering lysosomal function with GPN or the TPC2 agonist TPC2-A1-N protects zebrafish hair cells against delayed damage provide additional support to the idea that the lysosome is a critical regulator of hair cell death in this experimental system. These two agents result in release of calcium from lysosomes and an alkalinization in lysosomal pH^57,58,60^. TPC2 channels have also been shown to modulate iron levels and under some conditions TPC2-mediated iron release results in increase cell death through ferroptosis ^87^. However, our experiments suggest that TPC2 channels are not directly involved with the lysosome-mediated cell death process we observe after AG treatment. First, the TPC2 NAADP antagonist trans-Ned-19 alone has no effect on hair cell death, only reverses the protective effects of TPC2-A1-N. Second, we find that activating TPC2 is effective if we treat before AG exposure, suggesting that preconditioning alters the hair cell response to AG.

There are several potential mechanisms by which TPC2 activation acts to precondition lysosomes and protect against AG damage. Maintaining a critical pH balance inside lysosomes is crucial for the organelle catabolism functions, autophagic degradation, and macromolecule biogenesis ^88^. Moreover, changes in lysosomal pH have been shown to be a risk factor for diseases, like Alzheimer’s and Parkinson’s diseases ^89^. Lysosomal pH is a central regulator of iron metabolism ^67^. Cells with alkaline lysosomal pH can display an iron deficiency, as lysosomes are unable to convert the endocytosed Fe^3+^ into the Fe^2+^ form that is subsequently exported into the cytoplasm by the divalent metal transporter 1 (DMT1) ^73,90,91^. Lysosome alkalinization has been shown to increase resistance of senescent cells to ferroptosis ^66^.

An alternative mechanism would be that ion release promotes recruitment of lysosomal repair machinery that reduces permeabilization. Ca^2+^ signaling plays a central role in recruiting the ESCRT-III complex to lysosomes, where it serves to promote repair ^92^. Alternative lysosomal repair pathways such as ER-mediated repair regulated by PI4K2A ^93^ or repair through Annexin recruitment ^94^ are also activated through Ca^2+^ signaling. By contrast, lysosomal stabilization after damage through stress granule condensates is regulated by acidification ^95^. These different mechanisms likely interact with one another to cooperate in alleviating lysosomal damage ^96^. Under this scenario, treatment of hair cells with TPC2 agonist would potentially forestall AG-induced damage by poising repair machinery before AG exposure. Identifying mechanisms downstream of TPC2 activation may reveal potential therapeutic targets to prevent hair cell death.

Over the last decade the lysosome has emerged as an important signaling hub regulating metabolism, rather than just the catabolic center of the cell ^97^. Our studies support the idea that lysosomal disruption underlies some forms of AG-induced ototoxicity. Human mutations resulting in lysosomal dysfunction, in genes regulating lysosome maturation, homeostasis and metabolism including those causing lysosomal storage disorders have been shown to have profound effects on hearing, supporting the importance of this organelle for sensory hair cells ^98,99^. It will be interesting to determine whether genetic and ototoxic effects of lysosome perturbation share common elements.

## Ethics Statement

The animal study was approved by the University of Washington Institutional Animal Care and Use Committee. The study was conducted in accordance with the local legislation and institutional requirements.

## Author contributions

FBB: Conceptualization, Formal analysis, Investigation, Methodology, Validation, Visualization, Writing – review & editing, Funding acquisition. AC: Investigation, Formal analysis, Validation, Methodology. TL: Investigation, Methodology. DR: Conceptualization, Data curation, Funding acquisition, Project administration, Supervision, Writing – original draft, Writing – review & editing.

## Acknowledgments

We thank David White, the UW Zebrafish Facility for excellent fish care, and members of the Raible Lab for helpful discussion.

