## Supplemental material for "Lysosomal Ion Homeostasis Drives Delayed Hair Cell Death After Aminoglycoside Uptake"

**Figure S1**

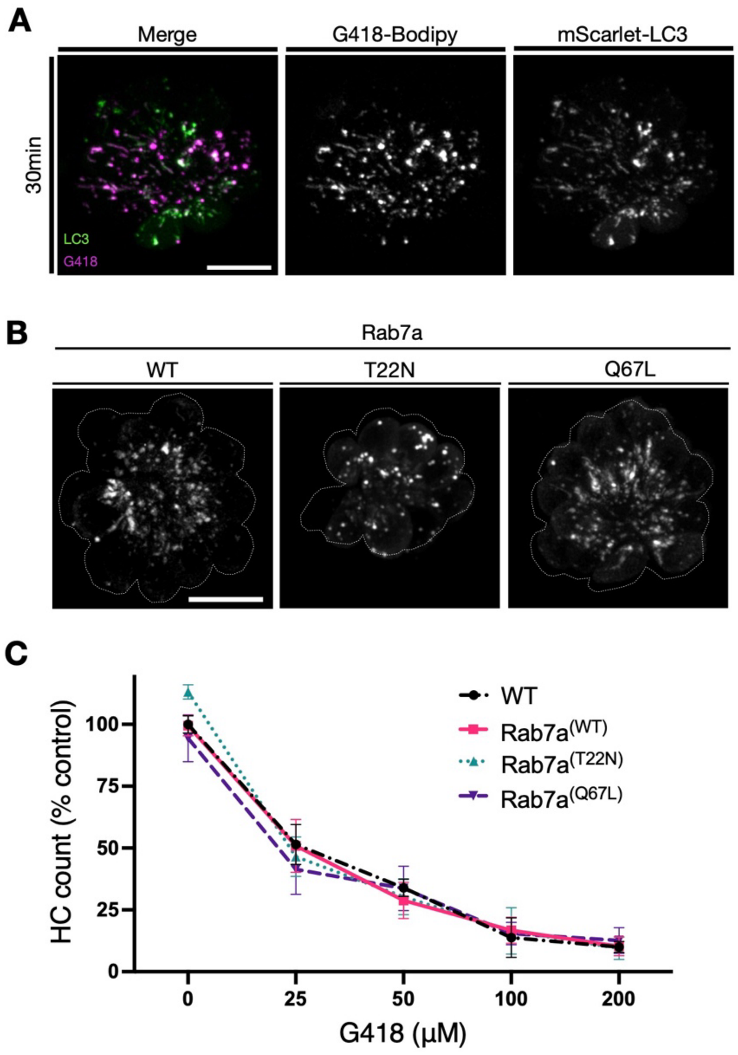

**Figure S2**

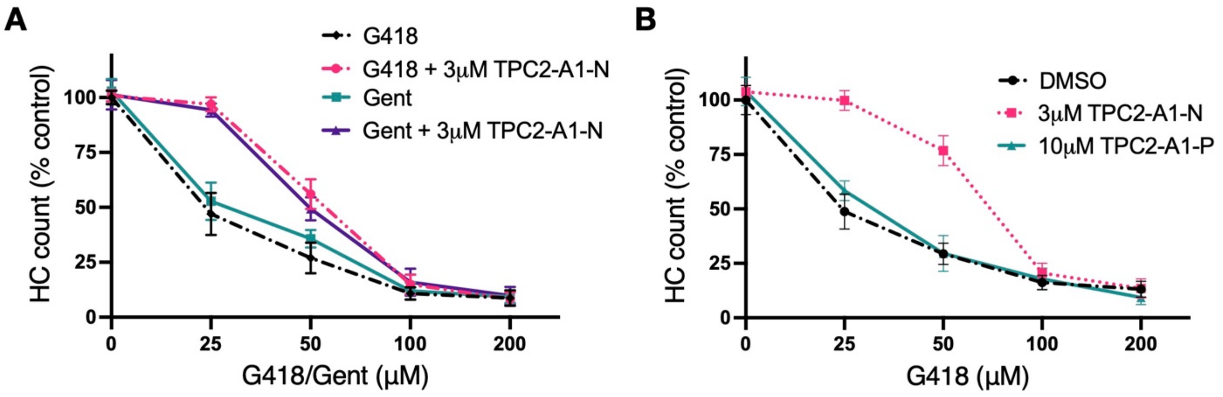
A
G418
G418 + 3μM TPC2-A1-N
Gent
Gent + 3μM TPC2-A1-N
HC count (% control)
100
75
50
25
0
0
25
50
100
200
G418/Gent (μM)
B
DMSO
3μM TPC2-A1-N
10μM TPC2-A1-P
HC count (% control)
100
75
50
25
0
0
25
50
100
200
G418 (μM)

**Figure S3**

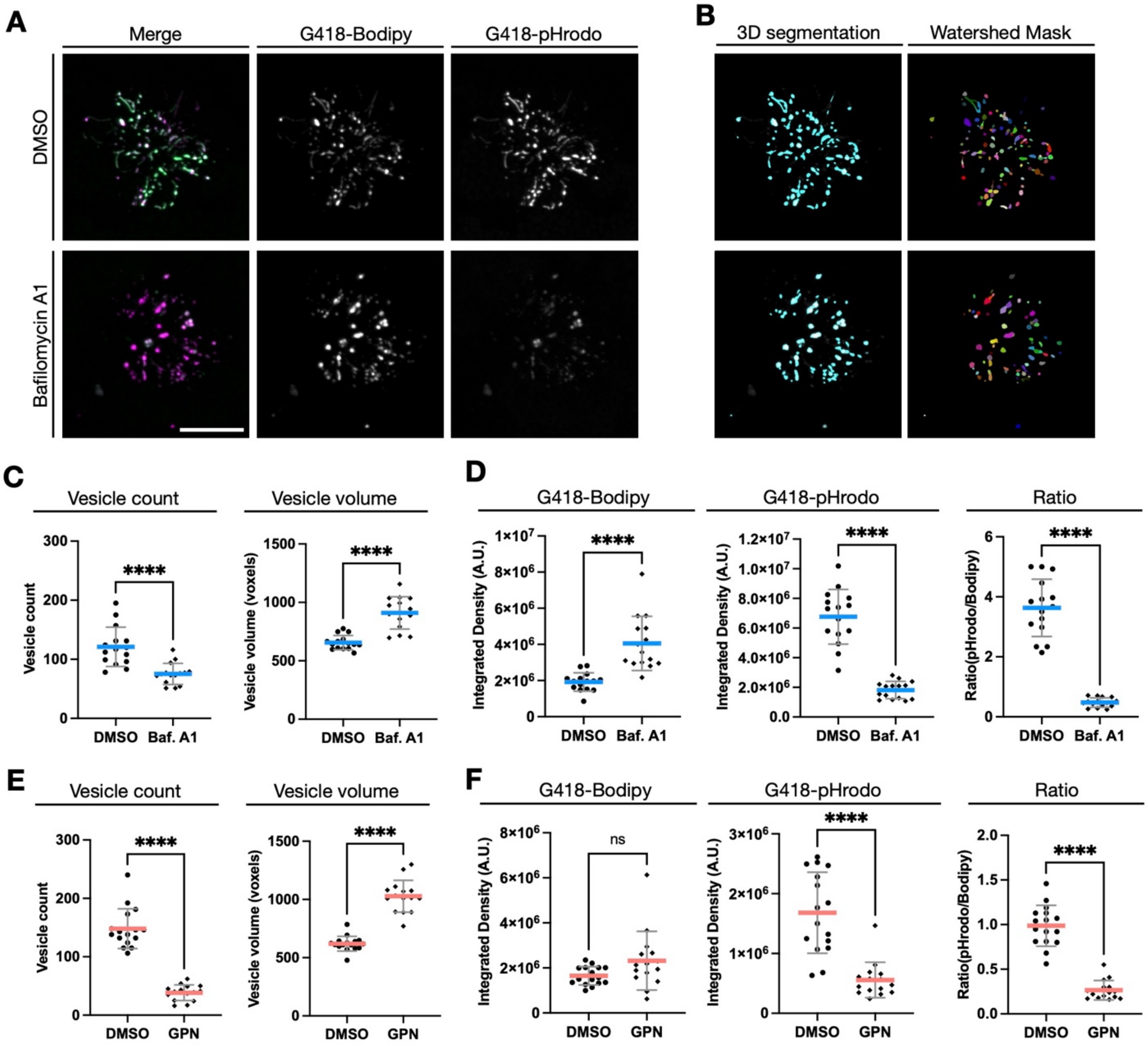

A
Merge
G418-Bodipy
G418-pHrodo
DMSO
Bafilomycin A1
B
3D segmentation
Watershed Mask
C
Vesicle count
Vesicle volume
****
Vesicle count
Vesicle volume (voxels)
DMSO
Baf. A1
D
G418-Bodipy
G418-pHrodo
Ratio
Integrated Density (A.U.)
Ratio(pHrodo/Bodipy)
E
Vesicle count
Vesicle volume
DMSO
GPN
F
G418-Bodipy
G418-pHrodo
Ratio
ns

**Figure S4**

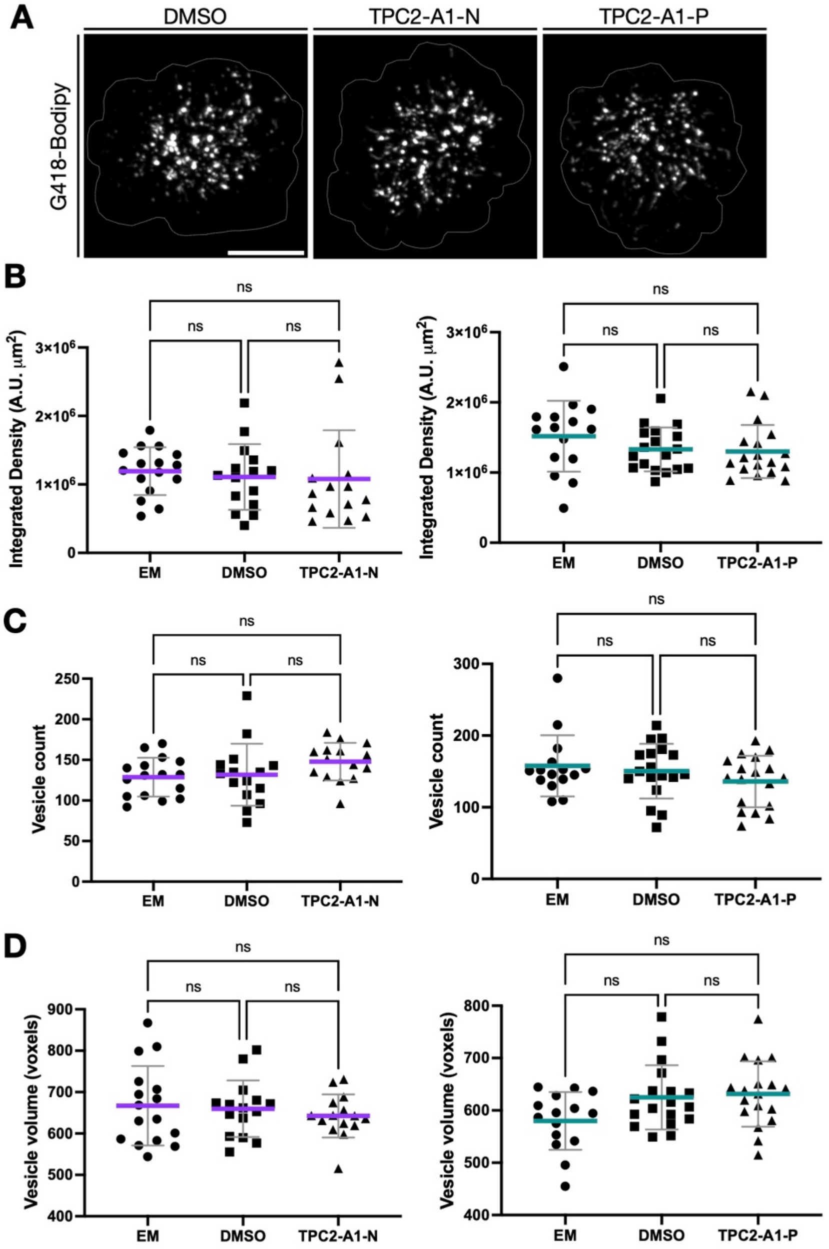

A
DMSO
TPC2-A1-N
TPC2-A1-P
G418-Bodipy
B
Integrated Density (A.U. μm²)
ns
EM
DMSO
TPC2-A1-N
TPC2-A1-P
C
Vesicle count
D
Vesicle volume (voxels)

**Supplemental Figure Legends**

**Figure S1: Rab7a mutations don't confer protection against delayed death. A)** 5dfp transgenic larvae, expressing mScarlet-LC3 (green) in hair cells, were pulsed with 50µM G418-Bodipy (magenta) for 30 minutes before imaging (Merge). G418 is being accumulated into LC3-positive vesicles. Scale bar: 10µm. **B)** Transgenic lines expressing mutated forms of mScarlet-Rab7a ($Rab7a^{(WT)}$) were created; dominant negative ($Rab7a^{(T22N)}$) and constitutively active ($Rab7a^{(Q67L)}$) forms. Expression of these forms visibly disrupt the morphology, localization, and number of Rab7a positive vesicles in hair cells. Dotted line delineates all hair cells in neuromast. Scale bar: 10µm. **C)** Wild type and transgenic larvae expressing different forms of Rab7a were exposed to G418 at different concentrations for 1h, washed, and incubated for 23h before counting surviving hair cells. Counts were normalized against EM control. No differences were found between the different groups. Stats: Two-way ANOVA, Individual comparison P-values: WT vs $Rab7a^{(WT)}$: 0.9902; WT vs $Rab7a^{(T22N)}$: 0.7165, WT vs $Rab7a^{(Q67L)}$: 0.4647.

**Figure S2: TPC2-A1-N but not TPC2-A1-P exposure protects against delayed hair cell. A)** 5dpf WT larvae were exposed to 3µM TPC2-A1-N for 1h, washout, and then exposed for 1h to G418 or gentamicin, larvae were then washed and incubated in EM for 23h. Surviving hair cell were counted, and counts were normalized against control. No differences were detected between G418 and gentamicin. Stats: Two-way ANOVA, Tukey's multiple comparison test P-value: G418 vs Gent: 0.4731; TPC2-A1-N G418 vs TPC2-A1-N Gent: 0.8247. **B)** 5dpf WT larvae were exposed to 3µM TPC2-A1-N or 10µM TPC2-A1-P for 1h, washed, and incubated for 1h to G418, larvae were then washed and incubated in EM for 23h. Surviving hair cell were counted, and counts were normalized against EM control. TPC2-A1-P is unable to protect hair cells against G418. Stats: Two-way ANOVA, Dunnett's multiple comparison test P-value: DMSO vs TPC2-A1-P: 0.3789; DMSO vs TPC2-A1-N: <0.0001.

**Figure S3: 3D segmentation allows for pH measurements in hair cell vesicles. A)** 5dpf WT larvae were incubated for 1h to 50µM of a mixture of equal parts of G418-Bodipy and G418-pHrodo green, washed and incubated 100nM Baf. A1 for 1h. Baf. A1 in the media was maintained during imaging. Representative images of neuromasts are shown (Merge). **B)** Using G418-Bodipy fluorescent signal vesicles were segmented. Single vesicles within the 3D mask where obtained after watershed processing. Single vesicle mask was then used to measure fluorescence intensity on both channels (Bodipy and pHrodo green), and mean vesicular ratiometric values were calculated. Size and number of vesicles were also calculated using mask information. **C)** Baf. A1 leads to a decrease of the average number of vesicles per neuromast, and an increase of their average volume. **D)** Baf. A1 exposure after G418 uptake leads to an increase average Bodipy integrated density, at the same time of a drop in pHrodo integrated density. Ratio calculation of integrated density shows an neutralization of the vesicular lumen. **E)** WT larvae were co-incubated with 250µM GPN and 50µM mixture G418-Bodipy and G418-pHrodo green for 1h. Larvae were then washed, but GPN in the media was maintained during imaging. Similarly to Baf. A1**,** GPN exposure leads to a decrease of the average number of vesicles per neuromast, and an increase of their average volume. **F)** GPN exposure during G418 uptake shows no change in Bodipy integrated density, at the same time of a drop in pHrodo integrated density. Ratio calculation of signals shows an neutralization of the vesicular lumen. Each dot corresponds to the average values of a single neuromast, 3 neuromasts per larva. Stats: Nonparametric Mann-Whitney t-test.

**Figure S4: TPC2-A1-N and TPC2-A1-P don't affect vesicle network. A)** WT larvae were preincubated with 3µM TPC2-A1-N or 10µM TPC2-A1-P for 1h. Drugs were then washed out and larvae were exposed to 50µM of a mixture of equal parts of G418-Bodipy and G418-pHrodo green for 1h. AG were washout before imaging. Images only depict G418-Bodipy channel. No important morphological changes in vesicles were observed between these conditions. Dotted line delineates all hair cells in neuromast. Scale bar: 10µm. **B)** No changes in G418 uptake were detected. **C-D)** 3D segmentation shows that after agonist exposure the number **(C)** and size **(D)** of vesicles inside hair cells remains unchanged. Stats: Kruskal-Wallis One-way ANOVA, with Dunn's multiple comparison test.
